# Genipin Alleviates Sleep Deficiencies Caused by α-Synuclein Toxicity in a *Drosophila melanogaster* Model of Parkinson’s Disease

**DOI:** 10.64898/2026.07.16.738995

**Authors:** Olivia M. Davis, Alice H. Sappenfield, Robert Fairman

## Abstract

Parkinson’s disease is predominantly characterized by dopaminergic neurodegeneration linked to toxic aggregation of α-synuclein. Genipin, a bioactive iridoid, was previously shown to improve the motility and survival deficits caused by pan-neuronal expression of native α-synuclein in a transgenic *Drosophila melanogaster* model system. We show that expression of α-synuclein causes sleep deficits and that genipin treatment rescued these sleep deficits, increasing total sleep and consolidating nighttime sleep relative to untreated α-synuclein–expressing fruit flies. Our findings extend genipin’s protective profile in *Drosophila melanogaster* and highlight sleep regulation as an additional phenotype responsive to α-synuclein–targeted interventions.

## Description

Parkinson’s disease (PD) is a dopaminergic neurodegenerative disease that manifests in the decline of motor function, tremors, rigid muscles, poor sleep, and loss of balance (Sjödahl et al., 2018). Complications from this disease typically lead to death as a result of the mass death of dopaminergic neurons (Ryu et al., 2023). This dopaminergic cell death is most commonly linked to the buildup of Lewy bodies, which include large-scale aggregates of protein α-synuclein (Wood-Kaczmar, Gandhi, Wood et al., 2006). *Drosophila melanogaster* serves as a valuable model organism to study transgenes, providing the ability to study human disease, including PD, through the expression of human α-synuclein. In flies expressing α-synuclein, both protein aggregation and behavioral effects are observed (Mizuno et al., 2010). A sleep study in *D. melanogaster* is warranted because of the high degree of conservation in the genes controlling circadian rhythm between humans and fruit flies (Dissel, 2020). Transgenic *D. melanogaster* expressing A30P α-synuclein has been shown to have fragmented and disrupted circadian rhythms, making this system both a useful model of PD and a target for therapeutic studies (Ito et al., 2017). Genipin, an iridoid found in various flowering plants, has been shown to have general anti-inflammatory effects and has been used in traditional medicine to treat central nervous system and inflammatory disorders (Koo et al., 2004). Studies in the primary literature have implicated genipin as a potential therapeutic for various neurodegenerative diseases, including rescuing developmental deficits in a mouse model of Familial Dysautonomia (Li et al., 2016; Saito-Diaz et al., 2023). In *Saccharomyces cerevisiae*, genipin has been shown to reduce α-synuclein aggregation and toxicity, although the disaggregase or inhibitory mechanism by which this occurs has not yet been clearly established (Rosado-Ramos et al., 2023). Additionally, genipin has been shown to improve mobility and survival in *D. melanogaster* expressing α-synuclein (Rosado-Ramos et al., 2023). Our study investigated the therapeutic potential of genipin within the context of the sleep deficits associated with PD.

Figure 1 shows the effect of α-synuclein expression on the sleep patterns of male *D. melanogaster* under a 12hr:12hr light:dark cycle and tests whether genipin can mitigate sleep disruption. We used a constitutively expressing neuronal driver line (elav-gal4) to activate α-synuclein expression in the experimental groups. The transgenic α-synuclein line was also crossed with the wild-type fruit fly (w1118) as a genetic control to account for the genetic background of the α-synuclein line. Sleep profiles of the flies were averaged over the three days of collection (Fig. 1A). Flies expressing α-synuclein slept more during the day (ZT 0-12) than the genetic control flies and appeared to have a lower sleep profile at night (ZT 12-24). These trends are consistent with those observed in previous work on mutated human α-synuclein (Ito et al., 2017; Xia et al., 2024). Numerical data from the sleep profiles were extracted and quantified for statistical analysis in Figs. 1B-D for ZT 12-24 and analyzed for potential mitigating effects of genipin.

**Figure 1.**
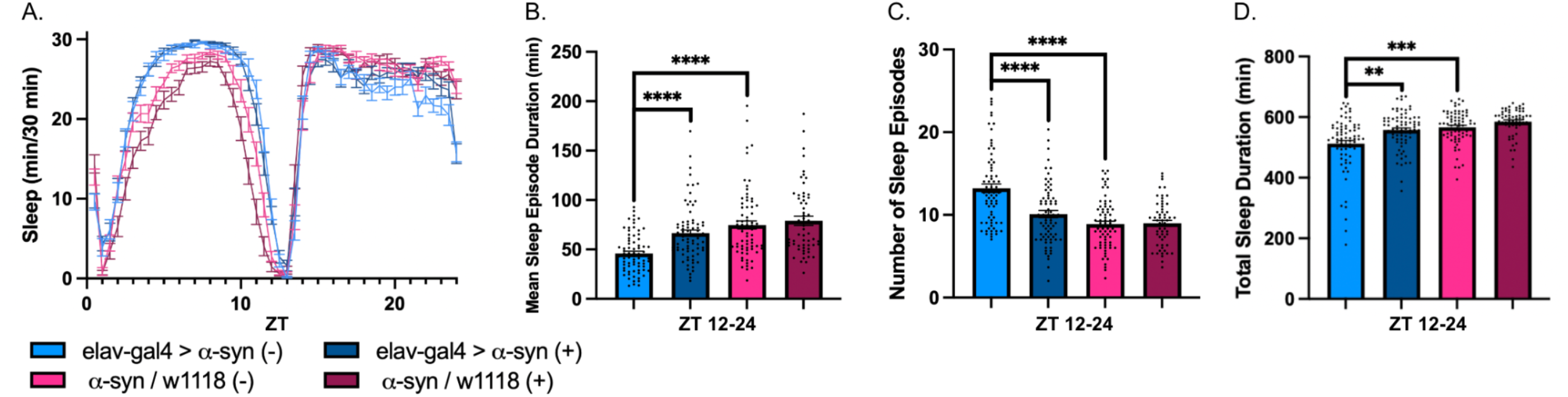
Genipin alleviates sleep deficiencies caused by α-synuclein expression in 7-10 day-old male fruit flies. Male flies expressing α-synuclein in the neurons were treated with (+) or without (-) 2 mM genipin for seven days post-eclosion and during the sleep study. Sleep data were analyzed over a period of three days in a 12hr:12hr light:dark cycle. The graphs show averages over the three-day bin. Treatment groups: elav-gal4 > α-syn (-) (n=74); elav-gal4 > α-syn (+) (n=76). Genetic controls: α-syn wt / w1118 (-) (n=72); α-syn wt / w1118 (+) (n=58). (A) sleep profile binned in 30-minute increments, (B) mean sleep episode duration in minutes, (C) number of sleep episodes, and (D) total sleep duration in minutes. These graphs are from three combined biological replicates and reflect the trends seen in each replicate. * P ≤ 0.05; ** P ≤ 0.01; *** P < 0.001; **** P ≤ 0.0001.

We first demonstrate that the constitutive pan-neuronal expression of human α-synuclein causes sleep deficits in *D. melanogaster* compared to their genetic controls. Flies expressing α-synuclein had a significantly lower mean sleep episode duration than that of the genetic control (Fig. 1B), a significantly higher number of sleep episodes compared to the genetic control (Fig. 1C), and a significantly shorter total sleep duration when compared to the genetic control (Fig. 1D). These combined graphs demonstrate that flies expressing wild-type human α-synuclein experience higher sleep fragmentation and shorter overall sleep. These findings justify testing whether genipin treatment can restore or reduce impaired sleep caused by α-synuclein expression.

We next demonstrate that genipin treatment rescues the sleep deficits caused by human α-synuclein expression. Flies expressing α-synuclein that were treated with genipin had a significantly higher mean sleep episode duration than that of their untreated counterpart (Fig. 1B). Genipin treatment restored the mean sleep episode duration of the α-synuclein-expressing flies to the wild-type phenotype seen in the genetic control. Treated α-synuclein expressing flies exhibited a significantly lower number of sleep episodes compared to untreated α-synuclein expressing flies (Fig. 1C). Combined with Fig. 1B, we demonstrate that genipin treatment reduces the sleep fragmentation caused by α-synuclein expression, restoring them to the wild-type phenotype observed in the genetic control flies. Additionally, flies expressing α-synuclein treated with genipin had a significantly higher total sleep duration at night than untreated flies expressing α-synuclein. Lack of significance between the treated and untreated genetic control groups indicates that we see no obvious general health effects of the genipin treatment, validating the impact of genipin on α-synuclein activity.

In this study, we showed that genipin treatment improves the sleep deficits caused by human α-synuclein expression in a *D. melanogaster* model of PD, proving to be a potential therapeutic for the sleep issues experienced by PD patients. Our findings suggest the possibility for future studies to investigate the molecular basis of how genipin interacts with α-synuclein pathology and, combined with the beneficial findings of previous work (Rosado-Ramos et al. 2023), suggest that genipin could serve as part of a future pharmacological therapy for patients with Parkinson’s disease.

## Materials and Methods

### Fly Husbandry

*D. melanogaster* stocks and experimental crosses were incubated at 25°C on standard cornmeal food media. Male progeny from experimental crosses were collected 0-2 days post eclosion and incubated at 25°C for 7 days with a 12hr:12hr light:dark cycle. Genipin (Biosynth) was dissolved in sterile Milli-Q water to a 20 mM stock solution and further diluted into sterile Milli-Q water to a 2 mM working concentration. It was orally supplied by adding it to standard fly media tubes for the 7-day incubation and incorporated into the sugar food medium for the sleep study. Experimental sleep studies were carried out in a 25°C incubator with a 12hr:12hr light:dark cycle.

### D. melanogaster Activity Monitoring and Data Analysis

Male fly progeny were placed into 65 mm Pyrex glass tubes (TriKinetics) individually with a sugar food medium (5% sucrose and 1% agarose [w/v], 2 mM genipin, sealed with wax) and confined using a cotton plug. Activity monitor tubes were placed into the *Drosophila* Activity Monitor (DAM) system and activity was monitored for 20 days using the Trikinetics DAMSystem3 program. Sleep data was analyzed using the Vecsey Sleep and Circadian Analysis MATLAB Program (SCAMP; v. R2024a) and averaged over 3 days. Statistical analysis was performed using GraphPad PRISM (v. 11.0.2) for statistical tests and graphing. Data that did not follow a normal distribution was analyzed using the Kruskal-Wallis test.

### Statistical Analytical Results

Statistical analysis results of sleep data, conducted on PRISM. A significance threshold of p < 0.05 was used for each test. All panels are from Figure 1.

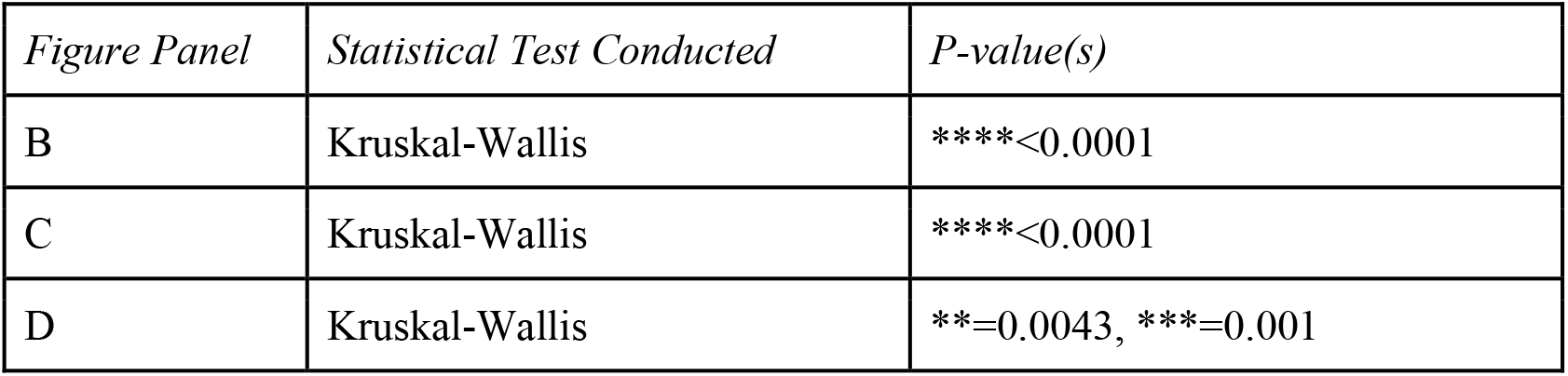

| Figure Panel | Statistical Test Conducted | P-value(s) |
| --- | --- | --- |
| B | Kruskal-Wallis | ****<0.0001 |
| C | Kruskal-Wallis | ****<0.0001 |
| D | Kruskal-Wallis | **=0.0043, ***=0.001 |

### Reagents

#### *D. melanogaster* lines used in sleep study

Reagents *D. melanogaster* lines used in sleep study.
| Fly Lines | Genotype | Source | Identifier |
| --- | --- | --- | --- |
| α-synuclein wt/sb | +/+; UAS-α-syn wt-eGFP/Tm6B,Hu,Tb | Domingos Lab, Lisbon, Portugal | - |
| elav-gal4/CyO | (Gal4-Elav.L)2/CyO | Bloomington Drosophila Stock Center | BDSC_8765 |
| w1118 | w[1118] | Bloomington Drosophila Stock Center | FBst0005905 |

## Acknowledgements

We thank Genevieve Uy and Dr. Pedro Domingos and his lab, for advice on this project. We thank Nicole Cunningham for technical assistance. Fly stocks were obtained from the Bloomington Drosophila Stock Center (NIH P40OD018537) and the Domingos Lab.

